# Coordinated electrical and chemical signaling between two neurons orchestrates switching of motor states

**DOI:** 10.1101/2023.01.04.522780

**Authors:** Maximilian Bach, Amelie Bergs, Ben Mulcahy, Mei Zhen, Alexander Gottschalk

## Abstract

To survive in a complex environment, animals must respond to external cues, e.g., to escape threats or to navigate towards favorable locations. Navigating requires transition between motor states, e.g. switching from forward to backward movement. Here, we investigated how two classes of interneurons, RIS and RIM, fine-tune this transition in the nematode *C. elegans*. By Ca^2+^ imaging in freely moving animals, we found that RIS gets active slightly before RIM and likely biases decision-making towards a reversal. In animals lacking RIS, we observed lowered Ca^2+^-levels in RIM prior to a reversal. Combined photo-stimulation and voltage imaging revealed that FLP-11, a neuropeptide released by RIS, has an excitatory effect on RIM, while tyramine, released from RIM, inhibits RIS. Voltage imaging of intrinsic activity provided evidence for tight electrical coupling between RIS and RIM via gap junctions harboring UNC-7 innexins. Asymmetric junctional current flow was observed from RIS to RIM, and vice versa. We propose that the interplay of RIS and RIM is based on concerted electrical and chemical signaling, with a fast junctional current exchange early during the transition from forward to backward movement, followed by chemical signaling, likely during reversal execution.

## Introduction

Throughout the animal kingdom, switching of behavioral states is indispensable to ensure survival and reproduction. During locomotion, animals generate appropriate motor patterns to execute complex movements. These include turning of direction, or switching from forward to backward movement. The latter requires active slowing, followed by transient stopping to then induce reverse locomotion. In vertebrates, the mesencephalic locomotor region (MLR) comprising the cuneiform (CnF) and the pedunculopontine (PPN) nuclei drive locomotion [1-3]. These two regions harbor glutamatergic neurons responsible for locomotion initiation as well as for speed regulation [4-6]. Furthermore, the PPN can induce locomotion stop via a rostral subpopulation of glutamatergic neurons [7], while GABAergic signaling from CnF and PPN fully inhibits locomotion [4, 5]. Next to the MLR, the gigantocellular nucleus (Gi) was shown to promote stopping behavior via glutamatergic V2a neurons [8, 9], depending on the strength of synaptic transmission: Strong unilateral stimulation of V2a neurons leads to stopping followed by turning [10]. In contrast, moderate activation results in a speed reduction prior to turning [11].

Neural populations mediating stopping or slowing during execution of directional changes appear to be a conserved feature present in evolutionary old vertebrate species like lampreys [12, 13] but also in invertebrates like *Drosophila* [14]. However, the details of how these stop neurons interact with other neurons to switch motor states remain largely unknown: Which chemical signaling is employed, how does electrical signaling contribute to induce stopping or slowing, and how do they precisely time the transition between forward and backward locomotion?

In the nematode *Caenorhabditis elegans*, these behaviors have been studied extensively. Sparse and whole-brain Ca^2+^ imaging experiments identified interneurons and premotor interneurons participating in regulation of forward (RIB, AVB, AIY) and backward (AVA/AVE/AVD, AIB, RIM) movement, some of which also orchestrate turns (AIB, RIB, AIY) [15-18]. Among the backward locomotion promoting neurons, AVA was shown to be instructive for reversal onset [19-22] and recently, [23] showed, that the second layer interneuron RIM represents a crucial hub for neuronal dynamics mediating the forward-reversal-turn transition. In a previous study, we found that the neuron RIS, which is not directly linked to AVA but shares chemical as well as electrical synapses with RIM [24, 25], is instructive for locomotion slowing and stopping. RIS achieves this by release of GABA, as well as FLP-11 neuropeptides [26]. This does not lead to paralysis, but the muscle tone is maintained, to enable quick resumption of locomotion. Thus, RIS regulates the early phase of switching motor states, and may be important for the precise timing of the constituent events. RIS also functions during sleep associated with developmental molting and sleep associated with starvation or satiety during which animals enter extended activity bouts, contrasting the brief events during locomotion control [27, 28]. In the latter context, we found compartmentalized Ca^2+^ dynamics in RIS: activity in the nerve ring (NR) was correlated with slowing of movement, while activity in a branch contacting the ventral nerve cord (VNC) was associated with reversals [26]. Interestingly, chemical synapses between RIS and RIM are restricted to the NR [25], while the branch contains mainly electrical synapses [24].

During reversals associated with an escape response, RIM inhibits the forward command interneuron AVB as well as head muscles via the release of tyramine [29, 30]. RIM is also involved in orchestrating omega turns [31, 32]. As shown by optogenetic depolarization, RIM can induce reversals; however, its ablation does not suppress them, and rather results in more frequent short reversals [20, 33, 34]. These differing properties of RIM may originate from its concomitant chemical and electrical signaling: RIM stabilizes forward ‘runs’ by acting as an electrical sink for AVA *via* gap junctions (GJs) containing UNC-9. Conversely, RIM uses chemical signaling to promote reversals when it is active [23]. Such combined signaling has also been observed for the coupling of AVA to A-type motor neurons to control (reversal) locomotion [22, 35], and appears to be a common feature during switching of motor states across species, especially during escape responses [36-39]. Chemical and electrical signaling influence each other directly. During development, GJs are required for the formation of intact chemical synapses [40-43] and chemical synapses play a role in the emergence and disappearance of GJs [44, 45], as well as in regulating junctional conductance [46, 47].

In sum, RIM employs dual signaling for sustaining reversals and shaping subsequent turning behavior [23, 31, 32]. Since it shares both chemical and electrical synapses with RIS [25], we wondered if and how RIM interacts with RIS in the early phase of reversal induction. During sleep behavior, ambiguous interactions of RIM and RIS were observed: Strong depolarization of RIM led to RIS inhibition, while moderate activation did not elicit a consistent response from RIS [48]. This may reflect a decision between a sole locomotion stop, or the induction of a subsequent reversal, which on the cellular level is reflected by the compartmentalized Ca^2+^-dynamics in RIS [26].

Here, we show that RIS and RIM orchestrate the chronology during reversal induction by the concerted use of chemical and electrical signaling. In freely moving animals, RIS became active before RIM, and in the absence of RIS, RIM exhibited a drop in Ca^2+^-levels prior to its activation and a reversal. Imaging of spontaneous voltage signals in immobilized *C. elegans* revealed bouts of fast periodic (up to 35 Hz) reciprocal electrical signaling. This suggests that RIS and RIM are tightly coupled through likely rectifying GJs that harbor UNC-7, and likely UNC-9, and the observed electrical activity may be a correlate for the delayed onset of RIM by RIS in moving animals. Photostimulation of RIS led to depolarized membrane potential in RIM via FLP-11 signaling, while depolarization of RIM demonstrated an inhibitory effect of tyramine on RIS. These findings implicate that charge is initially conferred from RIS to RIM, likely via GJ, biasing locomotion towards reversal induction. FLP-11 release from RIS further enhances RIM activity, thus stabilizing the reversal motor state. Subsequently, when RIM Ca^2+^-levels reach a plateau, acute release of tyramine inhibits RIS to enable the execution of the reversal, i.e. backward movement. Our study provides new insights in the fine-tuning of neurons involved in the transition of motor states, and in how behavior is achieved by orchestrated electrical and chemical synaptic transmissions.

## Results

### Simultaneous recording of Ca^2+^ activity in RIS and RIM neurites in freely moving animals

The RIS neuron is active upon slowing as well as when a stop occurs, and optogenetic stimulation of RIS induces these behaviors [26]. Ca^2+^ activity of the ventral branch of the RIS axon differed from the rest of the neurite, possibly as a result of local input. Upon reversals, Ca^2+^ increased in this branch as well as in the neurite in the NR. Slowing without reversals was accompanied by activity of only the NR axon, while the branch was silent. We thus asked how RIS interacts with other neurons to enable reversals, and whether this may need specific input in the branch. Among all neurons anatomically connected to RIS (**Fig. 1A, S1**), the RIM neuron was a promising candidate, as it mediates reversals; both neurons form chemical and electrical synapses with each other. Early connectomics work showed that gap junctions are located in the RIS ventral branch, along with other synapses [24]. We used our recent reconstructions of *C. elegans* brain anatomy to revisit the RIM-RIS synaptic connections [25] (**Fig. 1B, C; S2**). Chemical synapses from RIS to RIM predominate and are found exclusively in the RIS axon, with clear left-right lateralization (**Fig. 1B**). Few chemical inputs are received by RIS from RIM. RIS-RIM gap junctions were found in the RIS axon as well as in the branch (**Fig. 1C**). Note that the annotation shown in **Fig. 1B, C** is a summary of synapse identifications (see **Fig. S2A** for examples of gap junctions and chemical synapses in the RIS NR branch) observed in several animals as well as in [24] (**Fig. S2B; Supplemental Table 1**). The localization of RIM-RIS GJs in the branch was in line with the idea of a concerted electrical-chemical synaptic activity during reversal onset. We thus wanted to image activity of both neurons concomitantly during locomotion.

**Figure 1.**
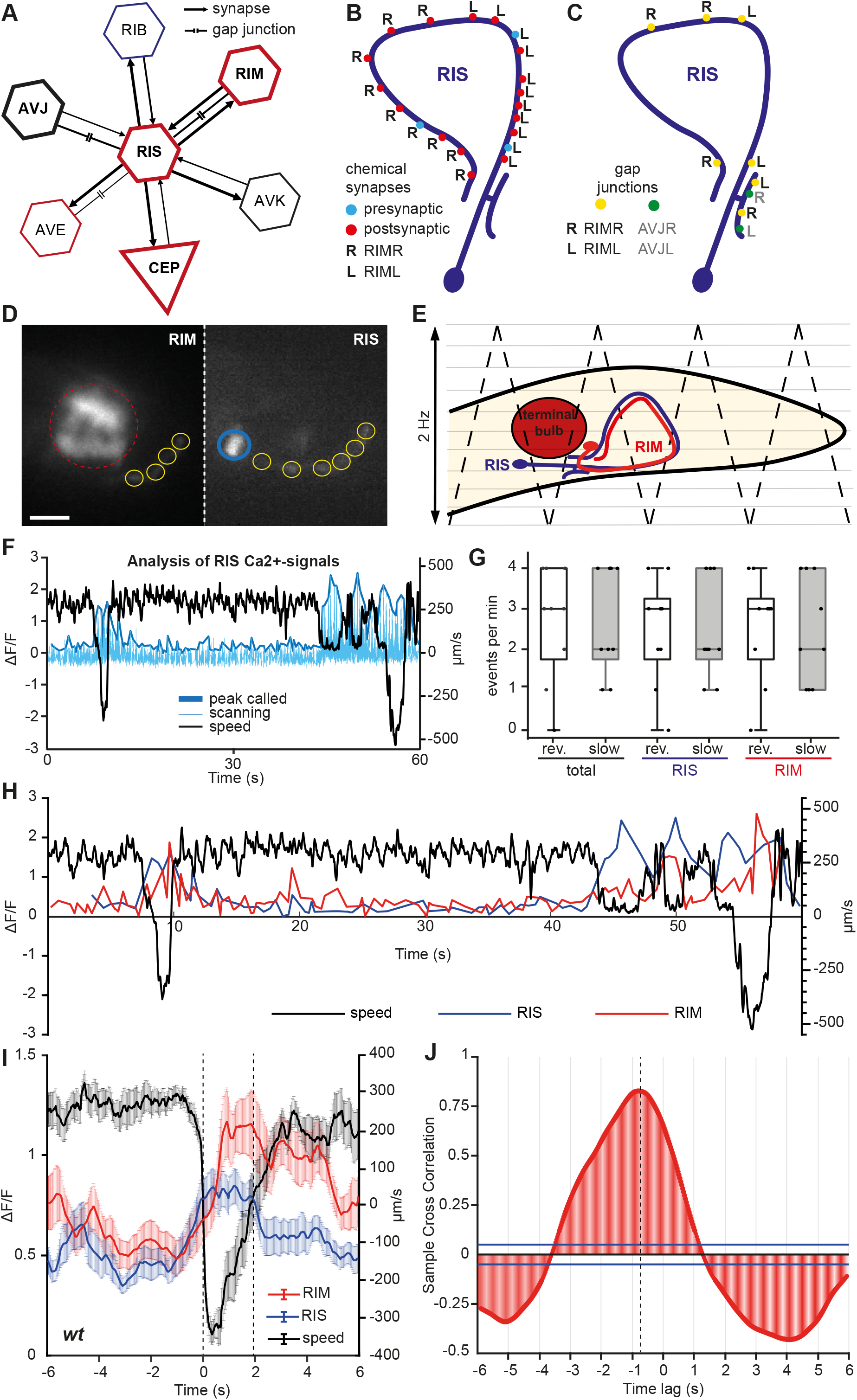
RIS and RIM axonal Ca^2+^-signals are coupled to reversals and temporally shifted. **(A)** Reduced connectome of RIS showing the pre- or postsynaptic neurons with the highest number of electrical or chemical synapses to RIS. Neurons associated with the reversal motor program or slowing response are marked in red, RIB, mediating forward locomotion speed, in blue. **(B)** Location of chemical or **(C)** electrical synapses between RIS and RIML/R cells, as indicated. **(D)** Fluorescence images from two different imaging channels used for tracking and Ca^2+^-imaging. Left: red channel used for visualizing jRCaMP1b expressed in RIM and tracking of a marker (mPlum) expressed in the pharyngeal terminal bulb (dashed red circle). Right: green channel used for visualizing GCaMP6s expression in RIS (blue circle: RIS cell body). Yellow circles indicate regions of interest (ROIs) alongside the axons of both neurons that are used for analyzing Ca^2+^-signals. Scale bar: 25 µm. **(E)** Schematic of z-stack acquisition from head region (dashed black lines indicate saw tooth scanning pattern). RIS (blue) and RIM (red) are located in different focal planes; terminal bulb used for tracking. **(F)** Peak-calling method for extracting Ca^2+^-signals of RIS. Raw values from z-scanning are background corrected and normalized (light blue). A custom-written MATLAB script extracts peaks (bold blue) and plots them along with the speed data (black). **(G)** Stopping and reversal events per minute, and fraction of these events during which activity of RIS and RIM was observed (slowing defined as speed <100 µm/s, activity of neurons assumed when Ca^2+^-signals increase by ≥30%). **(H)** Representative traces of velocity (dashed black line, scale on right y-axis), as well as RIS (blue) and RIM (red) neuronal activity (F/F_mean_, scale on left y-axis) monitored by GCaMP6s and jRCaMP1b signals respectively. **(I)** Animal velocity (black), Ca^2+^ signal amplitudes of RIS (blue) and RIM (red) aligned to locomotion stop during reversals (mean ± SEM; N=9 animals, n = 20 events); dashed lines at 0 and ∼2 sec indicate switching of forward to reverse, and again to forward locomotion, respectively. **(J)** RIS and RIM Ca^2+^-signals show significant cross-correlation (blue bars, p = ±0.05 confidence bounds) and a negative time lag of 840 ms (n = 20, Pearson’s r = 0.65).

As RIS and RIM are in different focal planes, we expanded our existing tracking system by the use of a Piezo objective scanner. Two spectrally distinct genetically encoded Ca^2+^ indicators (GECIs), GCaMP6s and jRCaMP1b, facilitated simultaneous axonal Ca^2+^-imaging in RIS and RIM (**Fig. 1D**). To this end, the head region of an animal, freely moving in the x,y-plane, was scanned in the z-dimension at approximately 2 Hz (**Fig. 1E**). Axons were tracked and Ca^2+^-signals in both imaging channels were extracted from regions of interest (ROIs) placed along the axons using TrackMate software in ImageJ (**Fig.1D**) [49, 50]. This method alleviates the need for image adjustment to accommodate distortion of the head morphology when imaging the axon. To find the focal plane of each neuron’s axon and to quantify its fluorescence, the raw fluorescent intensity data, as well as the position data obtained from the tracking stage were further processed with custom-written MATLAB scripts. These extracted the peak of the mean fluorescence of all ROIs from the fluorescence data stream that resulted from the oscillating objective focal planes (**Fig. 1F**).

### RIS activity precedes RIM activity during reversal induction

Confirming our previous findings, RIS activity coincided with slowing and reversal onset (**Fig. 1G, H**) [26]. RIM appeared to be most active during reversals, however, it also showed activity during slowing events, in line with previous findings (**Fig. 1G**) [23]. This could indicate an interplay between the two neurons.

To better understand the relative dynamics of RIS and RIM during the execution of reversals, we extracted 12 second time windows centered on the moment of zero velocity, when an animal executed a reversal, and averaged the Ca^2+^-levels of RIS and RIM as well as velocity (**Fig. 1I**). RIS Ca^2+^ rise coincided with the onset of slowing, which averaged about 1.5 second before the animal reached zero velocity, and attained a plateau along with the locomotion stop. Its activity lasted for about 2 s and started dropping during backward movement (**Fig. 1I**). Thus RIS likely supports slowing and initiation of reversals, but does not remain active for the entire duration of the reversal sequence. In contrast, RIM Ca^2+^-signals began to rise about 0.5 s after RIS and reached a plateau shortly after the maximal reversal velocity was reached. RIM activity dropped only after the animals resumed forward movement. Cross-correlation analysis of Ca^2+^ signals showed that RIS peak activity preceded RIM by about 0.8 s (**Fig. 1J**). RIS inhibits forward movement prior to a reversal, while RIM drives the reversal motor program. As their activity appears to be coordinated during reversal onset, this suggests an interplay of the two neurons that is required for the execution of reversals.

### RIM facilitates RIS activity by the release of tyramine

How might RIS and RIM interact? They are connected by both chemical and electrical synapses. Therefore, we tested these connections by analyzing mutants affecting each pathway. To facilitate reversals, RIM releases tyramine, which is considered to be an inhibitory transmitter [29-32, 51]. We thus compared the tyramine-deficient mutant *tdc-1(n3419)* to wild type, analyzing the Ca^2+^ activities of RIS and RIM relative to the reversal (**Fig. 2A**).

**Figure 2.**
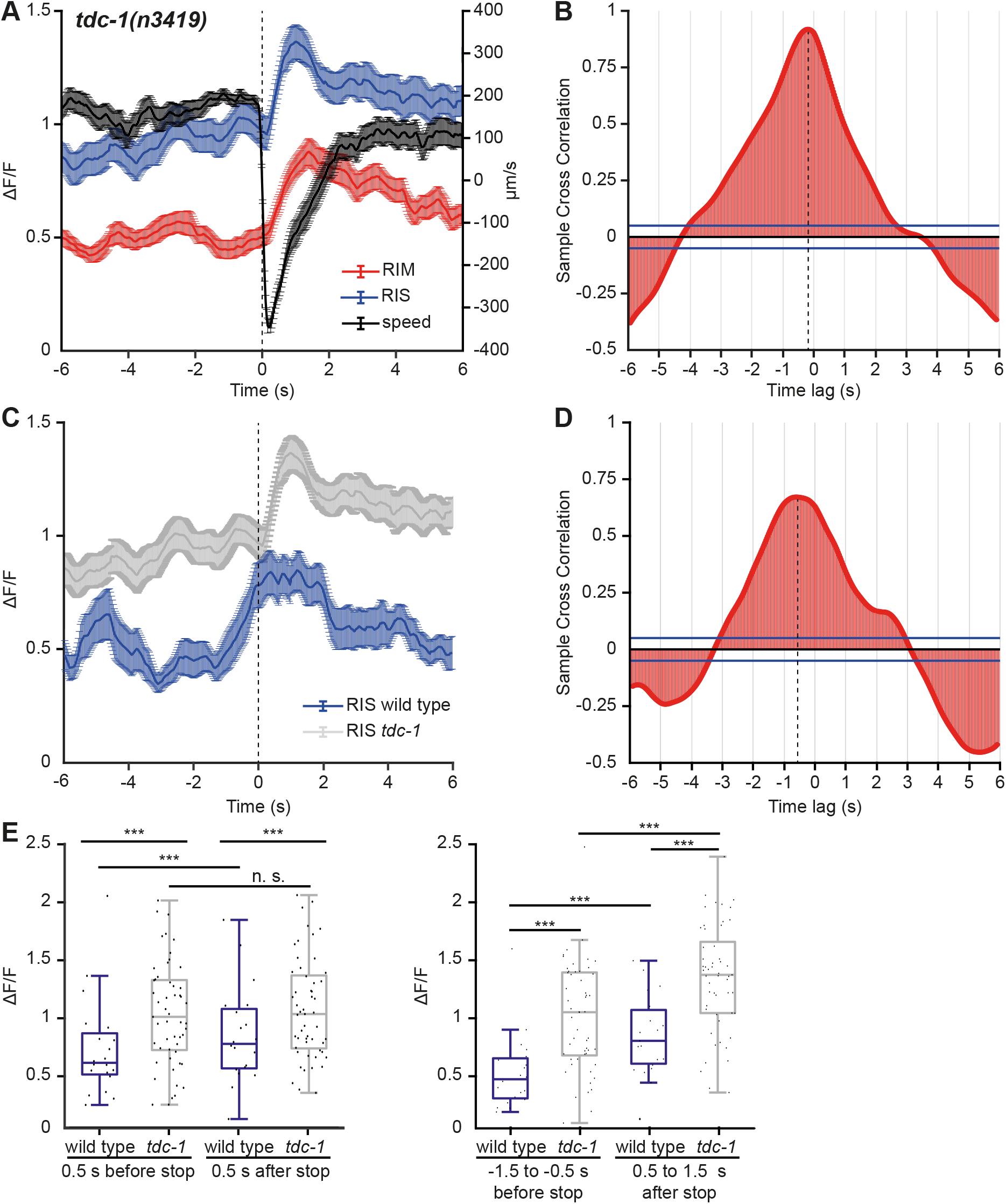
Tyramine is involved in regulation of RIS activity during reversals. **(A)** Animal velocity (black), Ca^2+^-signals of RIS (blue) and RIM (red) aligned to locomotion stop (dashed line) during reversals in *tdc-1(n3419)* background (mean ± SEM; N = 15 animals, n = 53 events). **(B)** RIS and RIM Ca^2+^-signals show significant cross-correlation (blue bars, p = ±0.05) and a negative time lag of 210 ms (Pearson’s r = 0.9). **(C)** Ca^2+^-signals of RIS in wt (blue, N = 9 animals, n=20 events) and *tdc-1(n3419)* mutant (grey; N = 15, n=53) aligned to locomotion stop. Note this data was acquired in parallel to data in **Fig. 1.** **(D)** Wild type and *tdc-1* Ca^2+^-signals in RIS show significant cross-correlation and a negative time lag of 570 ms (Pearson’s r = 0.63). **(E)** Analysis of Ca^2+^-levels in RIS during 1 s windows before and after locomotion stop in wild type and *tdc-1* background. Boxplot with Tukey whiskers. n = 53 events; *p ≤ 0.05; statistical significance tested by two-way ANOVA. In B and D, blue bars indicate 95% confidence bounds.

We found that the rise of RIS Ca^2+^-levels was delayed in *tdc-1* mutants and started only after reversal onset. Cross-correlation analysis revealed that RIS became active only 0.2 s prior to RIM (**Fig. 2B**), compared to 0.8 s in the wild type (**Fig. 1J**). Confirming the reduced delay between RIS and RIM in the *tdc-1* mutant, we found a delayed activity rise by ca. 0.6 s when we compared RIS Ca^2+^ levels in WT and *tdc-1* mutants (**Fig. 2D**). Since RIM activity onset was unaltered in the *tdc-1* mutant (**Fig. S3**), RIS must be delayed. The reduced delay in *tdc-*1 mutants may indicate a disinhibition of RIS in the absence of sustained tyramine levels, while acute tyramine release during the actual reversal leads to prolonged RIS activity. The RIS Ca^2+^ signal also exhibit altered peak amplitude and earlier decay in *tdc-1* mutants (**Fig. 2C**), consistent with RIS Ca^2+^ levels being significantly higher in *tdc-1* mutants both before and after locomotion stop (**Fig. 2C, E**).

In sum, findings in the *tdc-1* mutant indicate a role for RIM in promoting RIS activity, especially in the early phase of reversal induction and RIS activation. Given that tyramine is inhibitory, this may occur through disinhibition, or via an unknown tyramine-gated excitatory receptor in RIS.

### RIS promotes activity of RIM, preceding reversals

RIS was active prior to RIM during reversal induction, thus RIS may contribute to RIM activation. Previous work showed that RIM is stimulated by AVA and AVE, two premotor interneurons that are instructive for reversal locomotion. Yet, RIM can also negatively regulate reversals: RIM-ablated animals execute more brief reversals, and RIM activity drops during certain reversal events, while glutamate and tyramine signaling from RIM can both promote suppression of spontaneous reversals and increase reversal length [17, 20, 23, 31, 33, 35, 52, 53]. To address a role of RIS in RIM activation, we ablated the RIS neuron by overexpression of the apoptosis factor EGL-1, and recorded RIM Ca^2+^-signals during reversals, aligned to the moment of zero velocity.

Before the onset of RIM activity, we observed an ongoing reduction of RIM Ca^2+^-levels in the absence of RIS, but not in its presence (**Fig. 3A, B**). RIM Ca^2+^-levels reached their plateau just after the maximal reversal velocity occurred, while the rising phase was delayed in the RIS-ablated animals. The occurrence of the RIM peak was not altered in the presence or absence of RIS (**Fig. 3C**). Our data indicate that RIM might be gradually hyperpolarized during forward movement, as observed previously [23], and that RIS prevents this. RIS may thus gradually contribute to RIM activation. Ca^2+^ levels 0.5 s before or after the stop event were not significant different with or without RIS; however, Ca^2+^ levels before and after the stop differed significantly in the RIS-ablated animals (**Fig. 3D**). RIS thus attenuates the rise of RIM activity prior to a reversal, in line with RIM being active during slowing events (**Fig. 1G**). In sum, RIS exerts a basal effect on rising RIM activity, mainly before reversal onset.

**Figure 3.**
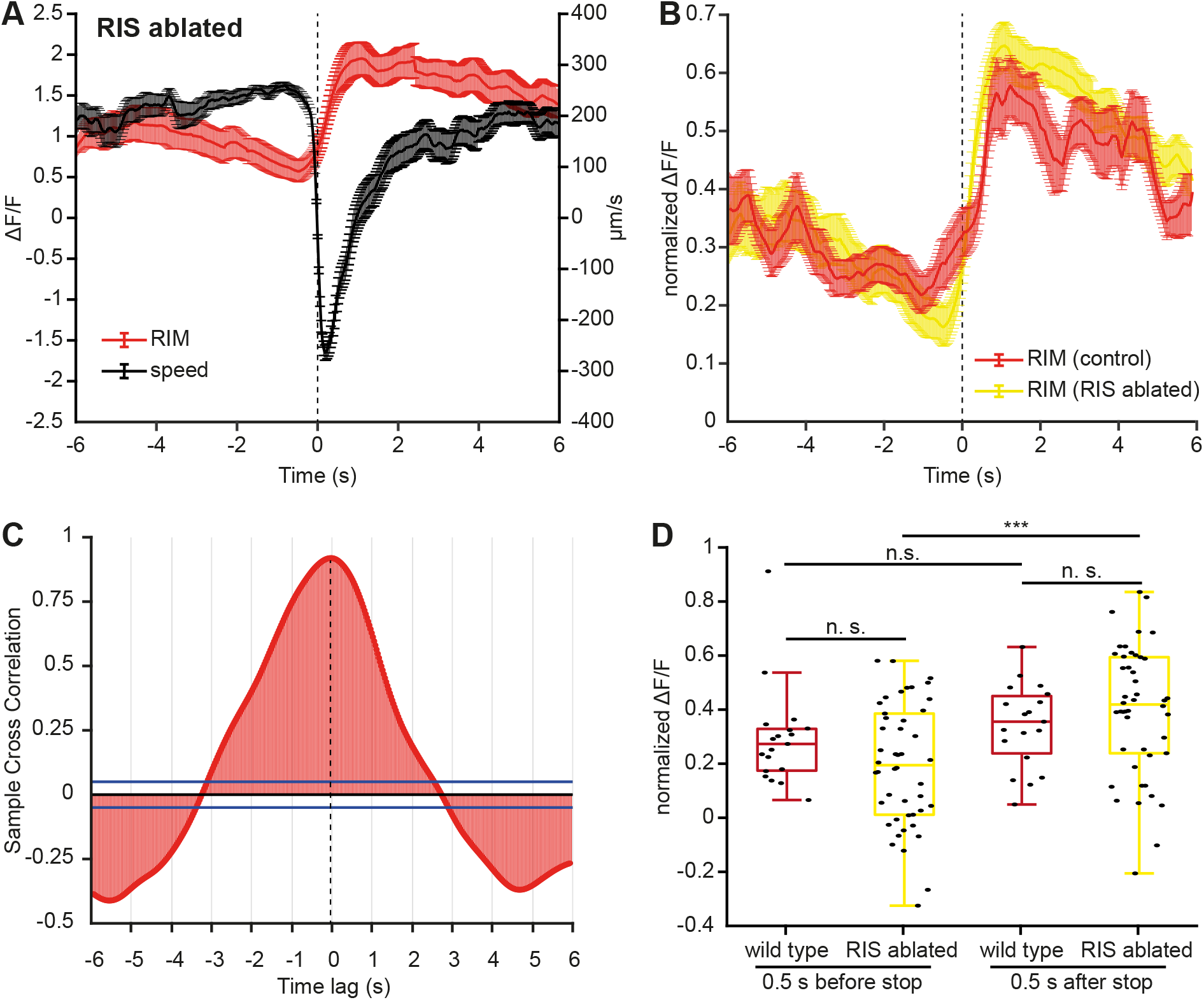
RIS promotes activity of RIM, prior to reversals. **(A)** Animal velocity (black), RIM Ca^2+^-signals (red), aligned to locomotion stop (dashed line, 0 s) during reversals in animals where RIS is genetically ablated (mean ± SEM, N = 15 animals, n = 47 events). **(B)** Normalized Ca^2+^-signals of RIM in wild type (red) and RIS-ablated animals (yellow), aligned to locomotion stop. **(C)** Normalized RIS Ca^2+^-signals in wild type and in RIS-ablated animals show significant cross-correlation and almost no time lag (-30 ms; n = 47 events, Pearson’s R = 0.92). Blue bars indicate 95% (p=±0.05) confidence bounds. **(D)** Analysis of normalized Ca^2+^-levels in RIS 0.5 s before and after locomotion stop in WT (N=9, n = 20) and RIS ablated animals (N = 15, n = 47) respectively. Boxplot with Tukey whiskers.; Significance tested by two-way ANOVA, ***: p<0.001.

### All-optical electrophysiology demonstrates functional coupling between RIS and RIM

Our analyses highlighted a likely interaction of RIS and RIM before and during initiation as well as execution of reversals. Ca^2+^ dynamics in the RIS axon are compartmentalized between its ventral axonal branch and the NR portion [26]. Possibly, gap junctions between RIM and RIS, located in its axonal branch (**Fig. 1C**) are involved in this activity pattern. Tyramine signaling may also play a role in RIM-RIS interactions (**Fig. 2B, C**). To address the complex interaction between RIS and RIM, but also between RIS and other neurons (**Fig. 1A**), we analyzed the electrical properties of RIS, and its electrical and chemical synaptic connections, using presynaptic optogenetic depolarization and post-synaptic voltage imaging.

In addition to locomotion slowing, RIS also induces a stop of pharyngeal pumping [26]. Among synaptic partners of RIS (**Fig. 1B, C**) that may be mediators of these effects are AVJ neurons, which are required for sustained high-frequency pharyngeal pumping [54], and CEP sensory neurons, which are responsible for slowing when animals enter a bacterial lawn; [24, 25, 55]. We photoactivated AVJ or CEP *via* channelrhodopsin-2 (ChR2), and examined RIS by voltage imaging, using the voltage indicator QuasAr2, tagged with GFP [56] (**Fig. 4A**), before, during and after the photostimulus (**Fig. 4B, D**). Voltage imaging was done at the cell body of RIS, as the axonal signal was too dim. The GFP signal was used to correct for motion or focal plane artefacts while QuasAr2 was imaged by excitation with a 637 nm laser. QuasAr2 fluorescence shows a voltage-independent increase in response to blue light. Thus, a strain expressing QuasAr2::GFP only was used as a control.

**Figure 4.**
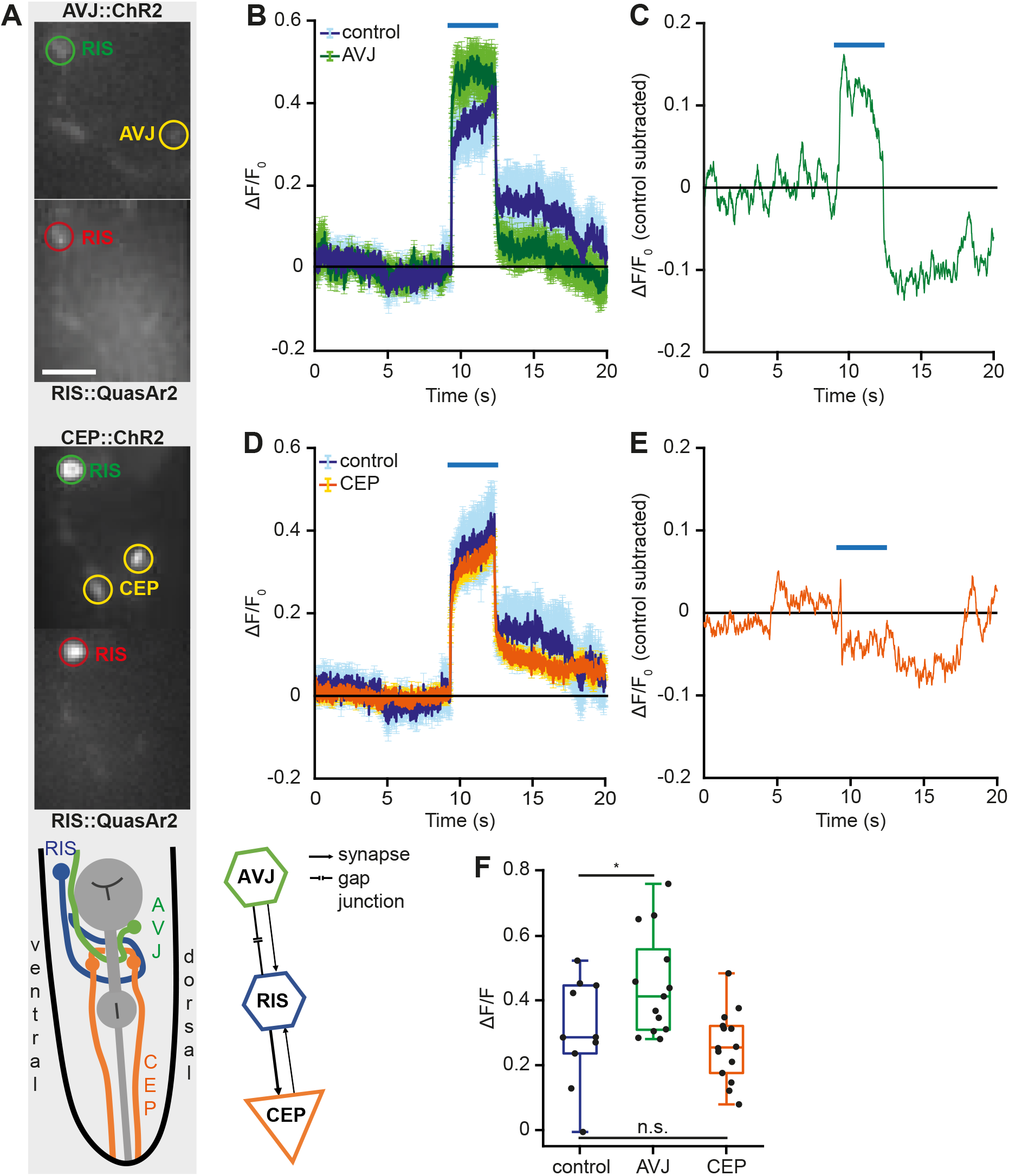
RIS voltage signals upon photo-stimulation of its synaptic partners AVJ and CEP. **(A)** Fluorescence images showing ChR2::YFP (yellow) and GFP (green) signals of AVJ and CEP, respectively, and red signal of the genetically encoded voltage sensor QuasAr2 in RIS. Scale bar: 25 µm. Lower panel: pictogram of the head, indicating position and orientation of RIS (blue), CEP (orange) and AVJ (green), pharynx in grey. Synaptic connections between these neurons are also shown in the scheme (lower right). **(B, D)** Mean (± SEM) voltage signals of RIS before, during and after 3 s blue light stimulation (blue bar) of animals with QuasAr2 expression only (ctrl, n=10) and animals expressing also ChR2 in AVJ (n=13), or CEP (n=14), respectively. **(C, E)** Difference in QuasAr2 signals between control group and animals with ChR2 expression in AVJ (green) or CEP (orange). Mean of control subtracted from mean of ChR2-expressing animals. **(F)** Analysis of voltage levels in RIS during the first 500 ms of photostimulation in controls (RIS::QuasAr2) and in animals additionally expressing ChR2 in AVJ (green), or CEP (orange), respectively. Boxplot with Tukey whiskers, statistical significance tested by one-way ANOVA (AVJ, * p = 0.048; CEP, p = 0.43 (n.s.)).

When we photoactivated AVJ, we observed a slight increase of QuasAr2 fluorescence in RIS, compared to the control strain (**Fig. 4B**). Subtracting the mean fluorescence intensities of the control from those obtained in the ChR2-expressing strain, revealed a rise of approximately 15% in ΔF/F_0_ during, and a drop of 10% after, blue light illumination (**Fig. 4C**). Comparing the voltage signals during the first 0.5 s, omitting the initial response to blue light (28 ms after stimulus onset), we found that the rise was significantly higher for AVJ photostimulation than in the control (**Fig. 4F**). This is in agreement with the idea that AVJ and RIS are electrically coupled and that depolarization of AVJ is transmitted to RIS. However, since AVJ also sends chemical synapses to RIS, using an unknown transmitter, the effects may also be due to excitatory chemical transmission. For photoactivation of CEP, we did not detect any obvious effects on RIS (**Fig. 4D-F**). This does not rule out that CEP may activate RIS, but given the low or variable number of synapses, this may simply be weak.

### RIS activates RIM via FLP-11 while RIM inhibits RIS via tyramine

Next, we conducted similar photostimulation and voltage imaging experiments in RIS and RIM (**Fig. 5A**). Photo-depolarization of RIM in wild type background led to neither an increase nor decrease of QuasAr fluorescence in RIS (**Fig. 5B**). This was unexpected, as there are both electrical and chemical synapses from RIM to RIS, and we previously found that tyramine signaling affected Ca^2+^ signals in RIS. We speculated that concomitant electrical (excitatory) and chemical (inhibitory) transmission may have canceled out in the optogenetic experiment. When we repeated the experiment in *tdc-1(n3419)* mutant animals, we observed a ∼15% relative increase in voltage signals compared to wild type animals (**Fig. 5B-D**). Thus, there is excitatory signaling from RIM to RIS, possibly through gap junctions (though we cannot rule out glutamatergic component), which is negatively regulated by tyramine.

**Figure 5.**
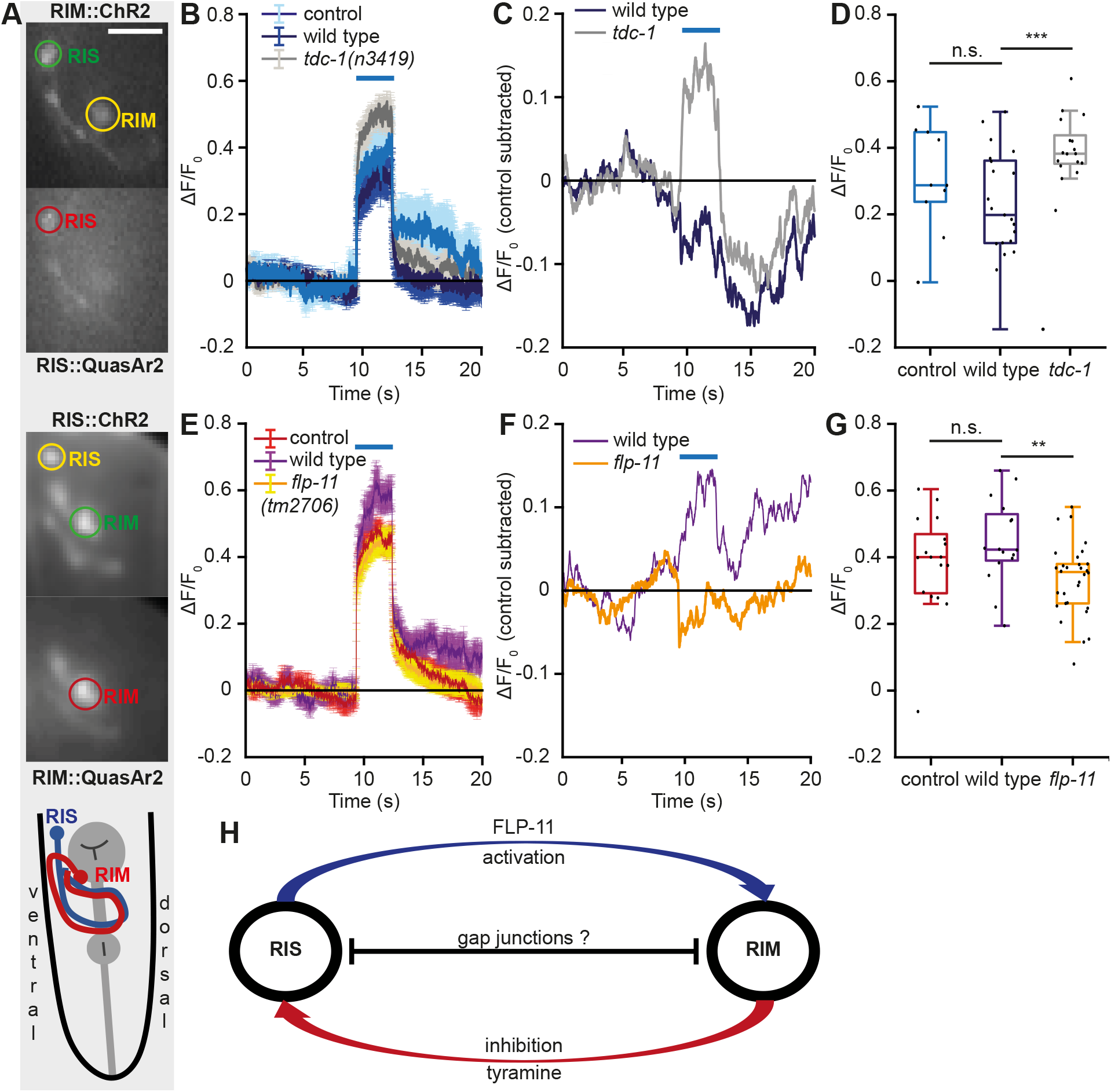
RIS and RIM voltage signals upon reciprocal photo-stimulation. **(A)** Top and middle: Fluorescence images showing expression of QuasAr2 (red) and ChR2::YFP in RIS and RIM, and vice versa. Upper panels: red channel (QuasAr2), lower panels, green channel (ChR2::YFP and GFP, tagged to QuasAr2), scale bar: 25 µm. Low: Pictogram illustrating position and orientation of RIS (blue) and RIM (red); pharynx in grey. **(B, E)** Mean (± SEM) voltage signals of RIS (B) and RIM (E) before, during and after 3 s blue light stimulation (blue bar) of animals with QuasAr2 expression only and animals expressing additionally ChR2 in RIM (ctrl: n=17, ChR2: n=21) and RIS (ctrl: n=10; ChR2: n=16), respectively. Also, photostimulation of RIM in tyramine-deficient *tdc-1(n3419)* mutants (n=17) and of RIS in *flp-11(tm2706)* neuropeptide mutants (n=30), respectively, was performed. **(C, F)** Difference of RIS::QuasAr2 signals between controls and animals with ChR2 expression in RIM, also for *tdc-1(n3419)* mutant animals (C), and of RIM::QuasAr2 in animals expressing RIS::ChR2 and control group, as well as in *flp-11(tm2706)* mutant animals (F). Mean of control group subtracted from mean of ChR2 expressing animals. **(D, G)** Statistical analysis of voltage levels in RIS during first 500 ms of photostimulation of wild type and animals expressing ChR2 in RIS or RIM, as well as in *flp-11* and *tdc-1* mutant animals, respectively, as indicated. Boxplot with Tukey whiskers, statistical significance tested by two-way ANOVA (C: p = 0,004; F: p = 0.003). **(H)** Model of RIS and RIM interplay. FLP-11 release from RIS excites RIM, while tyramine signaling from RIM inhibits RIS. Electrical synapses might be responsible for increased RIS voltage levels in the absence of tyramine.

In the reciprocal experiment, we observed an increase in RIM::QuasAr fluorescence upon RIS::ChR2 stimulation in wild type animals (**Fig. 5E, F**), demonstrating excitatory signaling from RIS to RIM. RIS uses the inhibitory transmitters GABA and FLP-11. Thus, this excitatory signaling may occur through RIS-RIM GJs. Yet, when we photo-depolarized RIS in the *flp-11(tm2706)* background, the resulting voltage levels were very similar to the control (**Fig. 5E, F**), while the voltage increase was significantly reduced compared to that of wild type animals (**Fig. 5G**). This was inconsistent with FLP-11 being an inhibitory transmitter. Possibly, *flp-11* release positively regulates RIM, e.g. through an unknown excitatory FLP-11 receptor, or FLP-11 neuropeptides may provide auto-inhibitory feedback to RIS itself. Taken together, we found that tyramine released by RIM acts inhibitory on RIS, while RIS positively regulates RIM *via* FLP-11 neuropeptidesor gap junctions. Furthermore, we suggest that electrical synapses are responsible for the activation of RIS upon RIM photostimulation in the absence of tyramine (**Fig. 5H**).

### RIS and RIM are reciprocally coupled via gap junctions harboring UNC-7 innexin

We found a complex interplay of RIM and RIS, mutually inhibiting or activating each other using chemical signals, but also *via* electrical synapses. In the absence of tyramine, RIM appeared to activate RIS *via* those gap junctions (**Fig. 5A-C**). We thus wondered about the nature of these GJs, depending on which, electrical coupling may serve to synchronize activity, or one cell can act as an electrical sink for the other. RIM expresses several gap junction subunits (UNC-7, -9, INX-1, -7 -18, and CHE-7, according to the RNA sequencing, www.cengen.org; [57]. Using such GJs, RIM stabilizes a hyperpolarized state of the reversal command neuron AVA during forward runs, and to regulate forward-to-reversal transitions [23]. RIS expresses INX-1, -7 and -14, according to CeNGEN; INX-1a, -1b, -2, -10a, -14, CHE-7, UNC-7, and UNC-9, according to [58]. UNC-7 and UNC-9 form heterotypic gap junctions [59, 60], thus connections of RIM and RIS should be affected in *unc-7(e7)* mutants.

To unravel details of inter-cellular signaling of RIS and RIM, we analyzed spontaneous voltage signals in animals expressing QuasAr2::GFP in both neurons (**Fig. 6A**). RIS and RIM both showed fluctuating activity, that often was not obviously synchronized, likely representing noise (about 4 % ΔF/F for RIM, and 7 % ΔF/F for RIS; **Fig. 6B**, upper panels, **Fig. S4A, B**). However, sometimes activity emerged significantly above noise level: This could be spiking activity, but also large fluctuations deviating from the basal level (-30 to +30 % ΔF/F for RIM, and +40 to -70 % ΔF/F for RIS; **Fig. 6B**, lower panels, **6C**). These large fluctuations were highly reciprocal in the two neurons, i.e. when RIS voltage went down, RIM voltage went up, and vice versa (**Fig. 6C**). Importantly, GFP signals obtained from both neurons showed no appreciable fluctuations and remained at noise level, less than 3 % ΔF/F (**Fig. 6B, C**).

**Figure 6.**
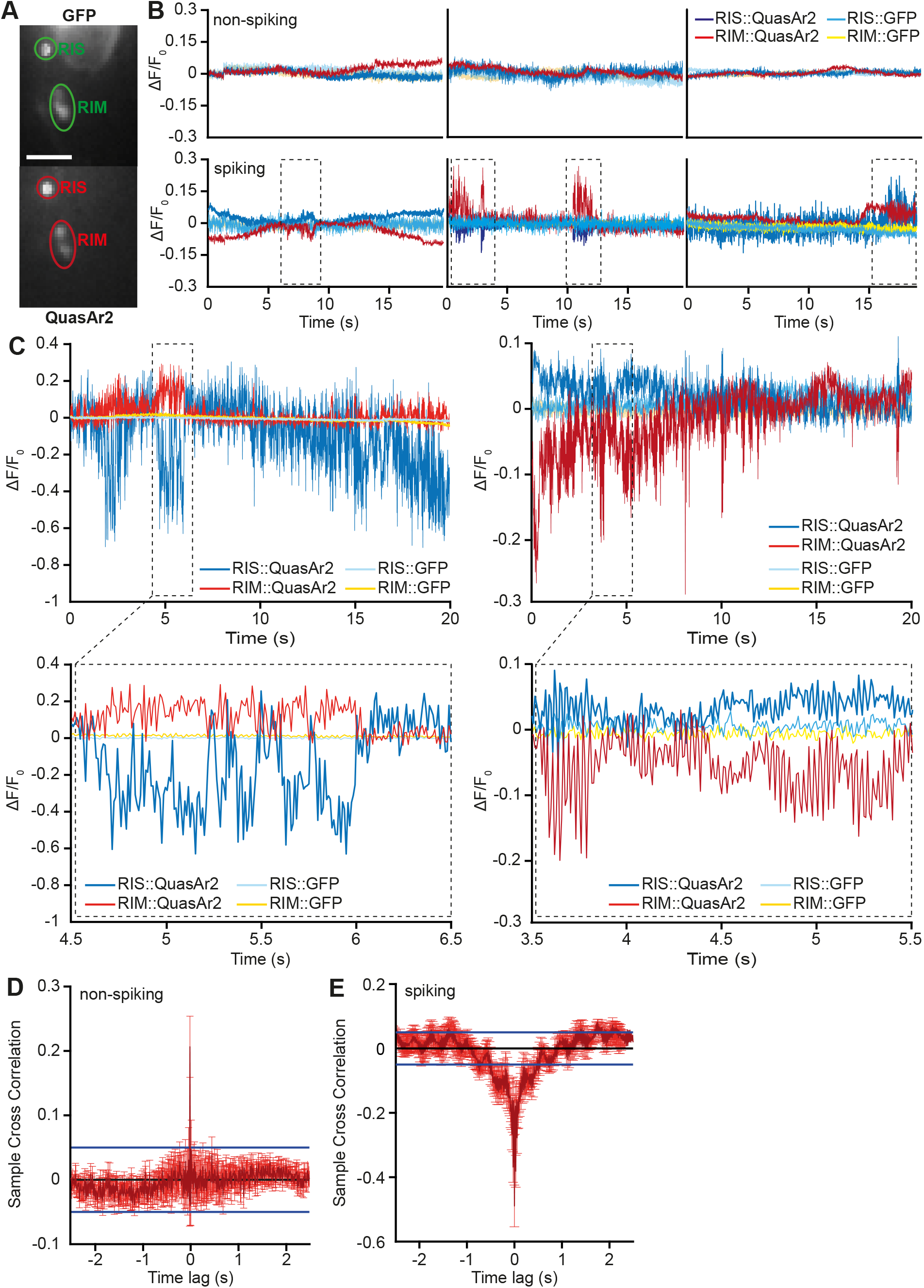
RIS and RIM exhibit reciprocal voltage signals. **(A)** Fluorescence images of dual voltage imaging experiments. Upper panel: signal of the GFP-tag. Lower panel: QuasAr signals of RIS and RIML/RIMR. Scale bar: 25 µm. **(B)** Representative voltage traces of RIS (blue) and RIM (red) imaging. Respective GFP signals of the QuasAr2::GFP fusion construct shown in light blue and yellow. Upper panels: example traces without spiking activity, lower panels: traces with spiking activity, indicated by dashed rectangles. **(C)** Upper panel: Enlarged traces of RIS (blue) and RIM (red) spontaneous voltage signals; examples for de- and hyperpolarizing episodes for both neurons. Control GFP signals shown in light blue and yellow. Dashed box refers to zoomed-in traces in lower panel, 2s windows with spike trains. **(D, E)** Cross correlation analysis of all voltage traces of RIS and RIM for the respective phenotypes. Mean ± SEM of 5 s time windows; blue lines indicate 95% confidence bounds (p=±0.05); Pearson’s r = 0.21 for non-spiking and 0.49 for spiking windows, each from N = 7 animals, n = 12 events. Spiking events were defined as activity exceeding noise levels, and not linked to movement (based on the GFP control).

We observed such reciprocal voltage events in about 10% of the recorded animals, probably because this spontaneous activity is rare upon immobilization. The events occurred in both directions: RIS exhibited positive and negative voltage fluctuations (de- or hyperpolarization) as did RIM (**Fig. 6B**, lower panels); however, in single animals, the signals of the respective neuron always deflected in the same direction, i.e. neurons would not switch from depolarized to hyperpolarized states. Possibly, it depends on an (unknown) internal state, which neuron is activated or inhibited. Of twelve event episodes we found for RIS, seven (five) exhibited an increase (decrease), respectively. In the same event bouts, for RIM, strictly reciprocal activity was observed. When we assessed the signals more closely, we found that even the smaller events, which were overlaid on the large fluctuations, i.e. single, spike-like activity, were reciprocal in the two connected neurons (**Fig. 6C**). These events were regular, occurring at up to 35 Hz. We assessed the extent of this coupling by cross-correlation analysis. While there was no obvious correlation when there was no activity (**Fig. 6D**), RIS and RIM were strongly anti-correlated (**Fig. 6E;** Pearson’s R = 0.49), with no detectable time lag. A single, small cross-correlation peak of non-spiking traces likely originates from camera noise, as it was also found when analyzing GFP signals of the same traces (**Fig. S4C**).

To assess whether electrical coupling (and activity) in RIS and RIM are GJ-dependent, we tested *unc-7(e7)* mutant animals in dual RIS-RIM voltage imaging. This innexin is not only present in RIM and RIS, and the mutant may thus be affected for other connections of these neurons that may contribute to the voltage events we see. Yet, imaging in these two neurons, and the correlation of signals in both cells, should primarily provide voltage information about RIS and RIM. These experiments showed much reduced spiking activity in RIS, and occasional spiking in RIM (**Fig. 7A**). Cross-correlation analysis of these signals, in 5 s time windows centered on the peak activity, exhibited almost no correlation at all (**Fig. 7B**). Fluctuations of the cross-correlation are probably due to the low number of events. Last, to examine whether the observed coupling of RIS and RIM voltage was based on chemical signaling, we analyzed spontaneous RIS and RIM signals in *flp-11(tm2706)* and *tdc-1(n3419)* mutants. The patterns we observed were very similar to those in wild type animals. Although the changes in ΔF/F_0_ appeared to be smaller (**Fig. 7C, E**), both mutants displayed the same strong anti-correlation with no temporal delay **(Fig. 7D, F)**.

**Figure 7.**
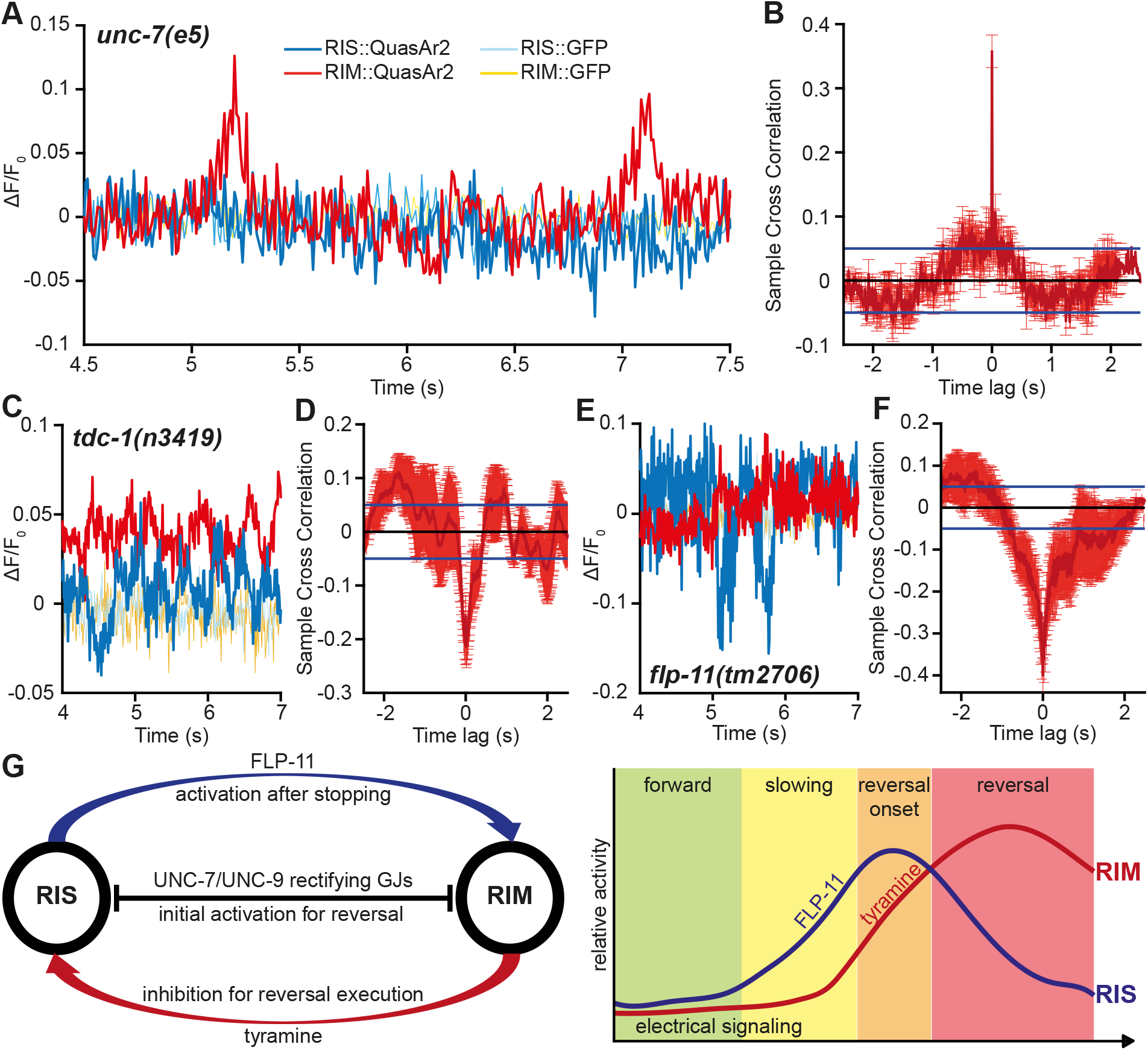
UNC-9/UNC-7 gap junctions mediate the tight electrical coupling of RIS and RIM. **(A, C, E)** Representative traces of voltage signals of RIS (blue) and RIM (red) in *unc-7(e5), tdc-1(n3419)* and *flp-11(tm2706)* mutant animals, respectively. GFP signals of QuasAr2::GFP fusion construct monitored in light blue and yellow; *unc-7*: N = 4 animals, n = 7 events; *tdc-1*: N = 3, n = 6; *flp-11*: N = 4, n = 7. **(B, D, F)** Cross-correlation analysis of the voltage traces in A, C, E of RIS and RIM for the respective genotypes. Mean ± SEM of 5 s time windows; blue lines indicate 95% confidence bounds (p=±0.05). **(G)** Model and time course of the interplay between RIS and RIM before and during a reversal. UNC-9/UNC-7 GJs mediate current flow from RIS to RIM and FLP-11 neuropeptides promote RIM activity after stopping. RIM in turn reduces RIS activity via tyramine signaling to enable the execution of the reversal motor program.

These findings demonstrate strong electrical coupling of RIM and RIS, with the negative cross-correlation most likely originating from GJs. The negative correlation might be explained by rectification. For example, during onset or execution of a reversal, positive charge may leave RIS (thus RIS membrane potential drops), and enter RIM (thus depolarizing it). The opposing event, i.e. where charge may enter RIS from RIM, thus depolarizing RIS and effecting a drop in RIM membrane potential, may possibly occur during quiescent states, or when the animal slows down without a reversal. The oscillations may be explained if the gap junctions would open only briefly, and opposing currents, such as tyramine- or glutamate-gated channels would revert the membrane potential back to the state before gap junction opening (**Fig. S4D**).

In sum, tight electrical coupling between RIS and RIM occurs *via* gap junctions harboring UNC-7, that may be selectively outward rectifying in RIS or RIM, and *vice versa*, depending on the state of the network of these neurons. Taking into account the Ca^2+^-imaging results from moving animals, we conclude that along with RIS activity onset, electrical coupling with RIM coordinates further events, such as release of FLP-11. This induces slowing and has an excitatory or disinhibitory effect on RIM, thus facilitating reversal induction. Upon rising RIM activity, it reduces RIS activity via tyramine signaling. RIM reaches highest activity when RIS Ca^2+^-signals start to decay. Jointly, these activities fine-tune, or even enable, execution of the reversal motor program (**Fig 7G**).

## Discussion

To overcome the challenges of navigating through a complex environment, vertebrates have evolved sophisticated neuronal systems, consisting of multi-layered circuits (Kiehn 2011). In the compressed nervous system of *C. elegans*, a relatively small population of neurons, the premotor interneurons, emerged to fulfill the basic tasks of locomotion, driving forward (AVB) and backward (AVA/AVE/AVD) movement. These command neurons interact with second layer interneurons such as RIS, RIM or RIB to fine-tune locomotion [19, 61, 62]. Of those, RIM is also able to induce reversals, while RIS mediates locomotion stop by the release of GABA and the neuropeptide FLP-11 [26]. RIS exhibits compartmentalized axonal Ca^2+^-dynamics, differing in the nerve ring and the ventral branch. Here, we showed how RIS interacts with RIM *via* concerted electrical and chemical signaling to orchestrate the chronology of steps during onset of a reversal.

RIS and RIM are tightly coupled via gap junctions that contain UNC-7, possibly with other innexins (**Fig. 6; 7A, B**). We suggest that these GJs are rectifying and can switch the polarity of rectification. Alternatively, gap junctions with both polarities are present and can switch from active to inactive states during different phases of the motor program. Different isoforms of UNC-7 have been shown to form heterotypic electrical synapses with UNC-9 that favor junctional current flow from UNC-9 to UNC-7 (UNC-7b) or *vice versa* (UNC-7e) in *Xenopus* oocytes [59].

Among interneurons driving backward locomotion, RIM is the only neuron presynaptic to RIS (**Fig. S1**). As RIS becomes active before RIM prior to reversal onset (**Fig. 1I, J**), it is plausible that junctional current flow occurs in the RIS to RIM direction to induce a reversal. The electrical synapses of RIS and RIM are, in part, located at the axonal branch of RIS (**Fig. 1C; S2**), which is instructive for reversal onset [26]. Rectifying GJs favoring outward currents would enable RIS to promote reversal induction, while its own activity is dampened, which may be required for reversals (**Fig. S4D**).

Our voltage imaging data was obtained in immobilized animals. However, it was previously observed that cyclic dynamics of cells in the *C. elegans* brain still occurs in immobilized animals [15]. We also observed events where RIS voltage increased, while RIM voltage dropped (**Fig. 6C**). This might reflect the events where slowing is not followed by a reversal, or possibly brief sleep states, when RIM activity needs to be dampened. How this is achieved by GJs remains unclear, as the composition of heteromeric UNC-7/UNC-9 GJs appears to be flexible [59].

The absolute voltage levels in both neurons cannot be deduced from voltage imaging. How the absolute voltage differs between RIS and RIM, as well as the dynamic range of each neuron, suggested by a higher range of ΔF/F in RIS than in RIM (**Fig. S4**) requires confirmation by electrophysiology. As this involves dissection, it might be difficult to correlate obtained data with events of behaviors.

During photo-stimulation of RIM, RIS exhibited inward currents in the absence of tyramine (**Fig. 5A-D**). This might originate from junctional current flow from RIM to RIS. However, chemical signaling of RIM besides tyramine could lead to elevated RIS voltage levels (**Fig. S4D**). Recently, RIM was shown to increase Ca^2+^ levels in muscle cells via the neuropeptide FLP-18 that binds to a GPCR, NPR-5, which is also expressed at low levels in RIS (www.cengen.org) [63]. RIM also releases glutamate, acting redundantly with tyramine to lengthen reversals [23, 64] and to excite RIS [48].

In moving worms, the onset of RIM activity followed that of the RIS NR axon, where GJs are prominent (**Fig. 1I**). Reversal induction is predominantly associated with fast activation of RIM by gap junction coupling with AVA [30], but before the animal reverses, it slows down forward locomotion [65]. Hence, it is possible that the fast activation of RIM by AVA is facilitated by RIS via UNC-7/UNC-9 gap junctions in the ventral branch of RIS. FLP-11 neuropeptides, released by RIS, are able to further enforce RIM activation [48]. Likewise, RIM activity (in the NR), and acute release of tyramine is subsequently needed for reversal execution.

RIM exhibited an increase of membrane voltage upon RIS photo-stimulation that was not observed in the *flp-11* mutant (**Fig. 5 E-G**). This indicates an excitatory role for FLP-11 in regulating RIM. As FLP-11 is considered to be inhibitory [66, 67], the excitatory effect of RIS on RIM could also be due to an indirect, dis-inhibitory effect, possibly *via* other neurons. However, during developmentally timed sleep, FLP-11 was shown to actively stimulate the forward pre-motor interneuron PVC [48]. This implies that an unknown excitatory FLP-11 receptor may also be expressed in RIM. Since PVC does not receive direct presynaptic input from RIS, this receptor would likely be a high affinity receptor, sensing FLP-11 in an endocrine fashion. Further, without RIS, RIM’s Ca^2+^ level was lowered prior to a reversal (**Fig. 3B, D**), while the timing of RIM activity was unaltered (**Fig. S3**). This is consistent with the idea of RIM being activated for reversal execution predominantly by AVA via GJs, but receiving electrical and chemical input from RIS to reinforce its activation and to further facilitate reversal induction.

Without tyramine, RIS’s basal Ca^2+^ level was increased (**Fig. 2A**) and RIM photo-stimulation resulted in higher voltage levels of RIS (**Fig. 5B**), in line with tyramine being inhibitory. However, RIS also had a smaller and shorter Ca^2+^-peak in *tdc-1* mutants (**Fig. 2C**), the onset of which was significantly delayed compared to wild type animals (**Fig. 2D**). The smaller peak might explain the phenotype of *tdc-1* mutants, exhibiting short reversals more frequently, with fewer long reversals [33]. As no excitatory tyramine-gated channel or receptor has yet been reported, disinhibition of RIS may promote this increased reversal activity. Basal tyramine levels might play a role in promoting RIS activity, probably by inhibiting neurons that normally inhibit RIS. In contrast, acute tyramine release during reversal execution could inhibit RIS, in agreement with the drop of Ca^2+^-levels in RIS when RIM becomes fully active (**Fig. 1I, 5A-D**). Inhibition of RIS is likely required for execution of the reversal and for subsequent steps, i.e. omega turns [33].

Here, we present a model where interactions between two neurons, RIS and RIM, fine-tune successive steps during the induction of the reversal motor program (**Fig. 7G**). Along with the onset of RIS activity, release of FLP-11 has an excitatory (or disinhibitory) effect on RIM, thus facilitating reversal induction. While RIM becomes active, it reduces RIS activity *via* tyramine signaling. During the early phase of activation of the backward command circuit, RIS induces locomotion slowing/stop and conducts junctional current to RIM via UNC-7 rectifying gap junctions, likely located in the ventral branch of the RIS axon. Subsequently, release of FLP-11 from the RIS NR axon leads to amplification of RIM depolarization, and stabilization of the reversal state. Finally, release of tyramine, presumably resulting from RIM activation by AVA, diminishes RIS activity, and enables the execution of reversals.

This principle - the same small subset of neurons are used for induction and inhibition of locomotion - is also found in the lamprey, where the MLR employs glutamatergic signaling to both elicit, and stop, a locomotor bout [12]. Also, the dual use of chemical and electrical synapses to tune neuronal output is found across species [37, 39]. Hence, the interplay between RIS and RIM might represent an ancient, conserved mechanism, integrating forward to backward transitions by the same subset of neurons.

## Acknowledgements

We are indebted to Franziska Baumbach for expert technical assistance and to Wagner Steuer Costa for kindly providing pWSC19. We thank members of the Gottschalk lab for critically reading the manuscript. Some strains were provided by the *Caenorhabditis* Genetics Center, which is funded by NIH Office of Research Infrastructure Programs (P40 OD010440). This work was supported by Deutsche Forschungsgemeinschaft (DFG) grant GO1011/13-1 to A.G., and by funds from Goethe University and the International Max Planck Research School in structure and function of biological membranes, to A.B. BM and MZ thank the past and current Zhen, Samuel and Lichtman lab members, the LTRI, and the CIHR (FDN154274) for supporting the connectome analyses presented in Figure S2 and Supplemental Table 1

## Author contributions

Conceptualization: M.B., A.G.

Methodology: M.B., B.M., A.B.

Software: M.B.

Validation: M.B., B.M., A.G.

Investigation: M.B., B.M.

Resources: M.B., A.B.

Data curation: M.B., B.M., A.G.

Writing – Original Draft: M.B.

Writing – Review & Editing: M.B., B.M, M.Z., A.G.

Visualization: M.B., B.M., A.G.

Supervision: M.Z., A.G.

Project Administration: A.G.

Funding Acquisition: M.Z., A.G.

## STAR Methods

### Molecular biology

Plasmid **18AALAOP** (18AALAOP_jRCaMP1b_pMA-RQ): codon optimized version of jRCaMP1b synthesized by Invitrogen (Thermo Fisher). **RM#348p** (*punc-17* vector) was a kind gift by James Rand. To generate **pXY09** (punc-17::jRCaMP1b), RM#348p was cut with KpnI and EcoRV and ligated to 18AALAOP cut with KpnI and EcoRI (blunted). **pBS77** (psto-3::GCaMP3) was provided by Zhaoyu Li (Xu lab, University of Michigan, USA). **pWSC15** (pggr-1::GFP) and **pWSC24** (pggr-2::flox::ChR2(H134R)::SL2::GFP) were provided by Wagner Steuer-Costa [26]. **pdat-1::ChR2::YFP** was provided by Martin Brauner [68]. To generate **pMF02** (psto-3::GFP), pWSC15 was cut with EcoRI and MscI and ligated to pBS77 cut with EcoRI and SmaI. Generation of **pXY07** (ptdc-1s::GFP): pMF02 was linearized with SphI and AgeI and ligated to the PCR product of ptdc-1s from wild type genomic DNA (forward primer oXY10: TCATGCATGCATTTCTGTATGAGCCGCCCG and reverse primer oXY15: TTGGACCGGTTGGGCGGTCCTGAAAAATGC) also cut with SphI and AgeI. **mPlum-N1** was a gift from Michael Davidson (Addgene plasmid #54629). **pncx-10::mCherry** was provided as linear PCR product by Petrus van der Auwera [26]. **pCG03** (pggr-2::flox::GCaMP6::SL2::RFP) was provided by Caspar Glock [26]. To create **pXY19** (ptdc-1s::GCaMP6), pCG03 was cut with AgeI and EcoRI (blunted), and ligated to pXY07, cut with AgeI and BsmI (blunted). **pAB23** (ptdc-1s::QuasAr::GFP) was described in [69]. **15AAYOCP-1670471-flp11prom** and **15ABJ3NP_1706249_3utrflp1** were gifts from Jan Konietzka. Plasmid **pXY26** (pflp-11::GCaMP6::SL2::RFP::flp-11_UTR) was generated by Gibson assembly with HiFi DNA Assembly Master Mix (NEB), using a restriction digest of pXY19 backbone with EagI and HindIII, and PCR products of pflp-11 from 15AAYOCP-1670471-flp11prom using primers oXY28 (AACAACTTGGAAATGAAATATTTGTTTTTTTGAAGGATTTTTGTG) and oXY29 (GAGATCCCATTATTCAGTATGAACTGCAAAAAGTG), GCaMP6::SL2::RFP from pCG03 with primers oXY30 (ATACTGAATAATGGGATCTCATCATCATCATC) and oXY31 (CATATGATTTCTATTTATAAAGTTCATCCATTCCATTAAG) and flp-11_UTR from 15ABJ3NP_1706249_3utrflp11 using primers oXY32: (TTTATAAATAGAAATCATATGTTTTTCTCTCTCACAC) and oXY33: (TGGAGCTCCACCGCGGTGGCGTTTTTGATCTAATATTTGAAAAAAAACAG). Plasmid **pXY29** (phlh-34::GCaMP6) was cloned from PCR product of phlh-34 from wild type (N2 strain) genomic DNA, amplified with primers oXY41 (TCAGCTATTACGGTGGTGGC) and oXY42 (CATACCGGTTCTCAAGTGGTTATAAGTCAAGCG), cut with AgeI and ScaI and ligated to pXY19, cut with AgeI and HindIII (blunted).

The following plasmids were used for generating strains: **pXY10** (ptdc-1s::jRCaMP1b) was generated from pXY09, cut with SphI and NheI (restriction site blunted) and ligated to pXY07, cut with SphI and AgeI (blunted). **pXY12** (pdat-1::ChR2::mCherry::SL2::GFP): For this, pWSC24 was cut with SphI (blunted) and Bsu36I, and ligated to pdat-1::ChR2::YFP, cut with PvuII and Bsu36I. **pXY17** (pncx-10::GFP) was cloned from pXY07, cut with SphI and AgeI and ligated to the PCR product of pncx-10 from pncx-10::mCherry, using primers oXY24 (TCATGCATGCTACACAGTTGCAGAGGCGTTTAATCAGA) and oXY26 (TCATACCGGTTACCTGAAAAAGAAACAGTTGATAAGCGGGT), also cut with SphI and AgeI. **pXY20** (pncx-10::mPlum), generated from pXY17, cut with AgeI and EcoRI and ligated to mPlum-N1, cut with AgeI and MfeI. **pXY23** (pggr-2::loxp::GCaMP6) was cloned from pXY19, cut with SphI and AgeI, ligated to pCG03 (cut with SphI and XmaI). **pXY27** (ptdc-1s::GCaMP6::SL2::RFP) was the ligation product of pXY19 ang pCG03, both cut with SphI and EcoRV. **pXY28** (pflp-11::QuasAr::GFP::flp-11 UTR) was generated from the PCR product of pXY26 amplified with primers oXY36 (ATACAAATAGAAATCATATGTTTTTCTCTCTCACAC) and oXY37 (TACTTACCATTATTCAGTATGAACTGCAAAAAGTG), and the PCR product of pAB23, amplified with primers oXY38 (ATACTGAATAATGGTAAGTATCGCTCTGC) and oXY39 (CATATGATTTCTATTTGTATAGTTCATCCATGCC), using Gibson assembly with HiFi DNA Assembly Master Mix (NEB). **pXY31** (ptdc-1s::ChRH134RT159C::YFP): pXY19 was cut with SpeI and AgeI (blunted) and ligated to pdat-1::ChR2::YFP, cut with SpeI and BamHI (blunted). **pXY32** (phlh-34 ChR2(H134R, T159C)::YFP): pXY29 was cut with SpeI and AgeI (blunted) and ligated to pdat-1::ChR2::YFP cut with SpeI and BamHI (blunted). **pWSC19** (pggr-1::Cre) was provided by Wagner Steuer Costa [26].

### *C. elegans* strains

N2 (WT, Bristol strain), **ZX2645**: *tdc-1(n3419); zxEx1240[pncx-10::mPlum; ptdc-1s::RCaMP1b-opti:SL2:GFP; pggr-2::loxP::GCaMP6::SL2::RFP; pggr-1::Cre]*, **ZX2564**: *zxEx1992 [pncx-10::mPlum; ptdc-1s::GCaMP6s:SL2:RFP]; goeIs384[pflp-11::egl-1::SL2::mKate2::flp-11-3’-UTR; unc-119(+)]*, **ZX2867**: *unc-7(e5); zxIs129[ptdc-1s::QuasAr::GFP; pELT-2::GFP]; zxEx1381 [pflp-11::Quasar::GFP::flp-11-3’-UTR]*, **ZX3178**: *zxEx360 [pggr-1::Cre, pggr-2::flox::ChR2(H134R)::mCherry::SL2::GFP]; zxIs129[ptdc-1s::QuasAr::GFP; pELT-2::GFP]*, **ZX3179**: *flp-11(tm2706)X; zxEx360[pggr-1::Cre, pggr-2::flox::ChR2(H134R)::mCherry::SL2::GFP]; zxIS129[ptdc-1s::QuasAr::GFP; pELT-2::GFP]*, **ZX3180**: *zxEx1294[pflp-11::Quasar::GFP::flp-11-3’-UTR; phlh-34::ChR(H134R/T159C)::YFP]*, **ZX3181**: *zxEx1293[pflp-11::Quasar::GFP::flp-11-3’-UTR; ptdc-1::ChR(H134R/T159C)::YFP]*, **ZX3182**: *tdc-1(n3419); zxEx1293[pflp-11::Quasar::GFP::flp-11-3’-UTR; ptdc-1::ChR(H134R/T159C)::YFP]*, **ZX3183**: *zxIs129[ptdc-1s::QuasAr::GFP; pELT-2::GFP]; zxEx1381[pflp-11::Quasar::GFP::flp-11-3’-UTR]*, **ZX3184**: *tdc-1(n3419); zxIs129[ptdc-1s::QuasAr::GFP; pELT-2::GFP; zxEx1381[pflp-11::Quasar::GFP::flp-11-3’-UTR]*, **ZX3185**: *flp-11(tm2706)X; zxIs129[ptdc-1s::QuasAr::GFP; pELT-2::GFP]; zxEx1381pflp-11::Quasar::GFP::flp-11-3’-UTR]*, **ZX3186**: *zxEx1295 [pflp-11::Quasar::GFP::flp-11-3’-UTR]; pdat-1::ChR2(H134R/T159C)::YFP]*.

### *C. elegans* cultivation and transgenic animals

All strains were kept at 20 °C on nematode growth medium (NGM) plates seeded with *Escherichia coli* OP-50-1 bacteria. For photostimulation and/or voltage imaging experiments 100 µm all-trans-retinal (ATR, Sigma-Aldrich) was added to the bacterial culture used to seed the plates and L4-stage animals were transferred to the plates one day before the experiments. Transgenic animals were generated by microinjection and extra-chromosomal arrays were integrated by UV-light illumination following standard protocols.

### Ca^2+^-imaging setup

Our previously described setup (Steuer-Costa et al., 2019) was equipped with a PIFOC P721 objective scanner and an E-709.CRG Digital Piezo Controller (both Physik Instrumente (PI) GmbH & Co. KG, Germany).

### Ca^2+^-imaging in freely moving worms

Image acquisition was performed as described previously [26]. Only minor changes were applied. Expression of the terminal bulb marker was changed from pncx-10::mCherry to further red shifted pncx::mPlum to reduce the fluorescence of the bulb marker in the red imaging channel. Additionally, jRCaMP1b was expressed in RIM with the *tdc-1* promotor. Fluorescence of jRCaMP1b and mPlum was excited with a 100 W mercury lamp equipped with a 575/40 ET bandpass filter (F47-573, AHF) and KSL-70 LED lamp (Rapp OptoElectronic, Hamburg, Germany) was solely used for excitation of GCaMP6s. Z-scanning at 2 Hz with the Piezo objective scanner did not influence tracking as the terminal bulb structure spanned all focal planes.

### Image analysis and data processing

Image stacks were split into red and green channel using crop3D function and terminal bulb structure was cleared manually in ImageJ. TrackMate software in ImageJ was used to extract Ca^2+^-signals from both GCaMP6s in RIS and jRCaMP1b in RIM. Circular regions of interest (ROIs) were defined, capturing fluorescent structures alongside the axons. The diameter was chosen slightly larger than the actual fluorescence signals and non-fluorescent areas were used for background correction later on. Parameters were adjusted to allow only axonal fluorescence to be extracted, in some cases this required a manual proofread of the tracks. Raw data was saved to the same Excel file containing speed data from the xy-stage and sorted by frame numbers. Further processing was performed in Matlab (MATLAB R2018a, MathWorks, USA): (1) background correction was performed by subtracting the signal of the non-fluorescent areas from the actual fluorescence signals. (2) DF/F_mean_ was calculated. (3) The *findpeaks* function was used to extract the Ca^2+^-curves from the z-scanning data (Fig. 1e). (4) Speed data was smoothed by a moving average of window size k = 5 frames. (5) Ca^2+^-signals were also normalized to the maximum value. (6) results were saved to the same excel file and plotted in figure containing speed, RIS-Ca^2+^-, RIM-Ca^2+^-data.

### Photostimulation and voltage imaging in immobilized *C. elegans*

Animals were immobilized on 10% agarose pads with polystyrene beads (Polysciences, USA) and imaged with a 40x oil immersion objective at Zeiss Axio Observer (Zeiss, Germany). Quasar fluorescence was excited with a 637 nm red laser (OBIS FP 637LX, Coherent) at 1.8 W/mm2 and imaged at 700 nm (700/75 ET Bandpass filter, AHF). GFP and ChR2(H134R)) were excited/activated using a monochromator (Polychrome V, Till Photonics/Thermo Scientific) with a bandwidth of 10 nm at 1 mW/mm^2^ and 100 µW/ mm^2^ respectively. A 497/655 H dualband beamsplitter (F58-200, AHF) and DualView2 (Photometrics, USA) were used for dual channel imaging. GFP emission was imaged with a 532/18 Brightline HC bandpass filter (F39-833, AHF). For photostimulation, videos were cropped and synchronized to the blue light pulse using crop3D function in ImageJ [50]. Movements of the cell body were corrected using trackmate software in ImageJ [49]. Circular ROIs with mean QuasAr fluorescence values for the cell bodies (CBs) and the background (inside the tissue), which was subsequently subtracted, were defined. Afterwards, dF/F_0_ was calculated, where F_0_ is the mean fluorescence before blue light stimulation. The same was done for the signal of the GFP-tag, which served as control for movement or CB deformation. For dual voltage imaging dF/F_mean_ was used.

### Signal cross correlation analysis

Cross correlation analysis was performed in Matlab (MATLAB R2018a, MathWorks, USA) by calculating Pearson’s correlation functions for equally sized 6 s time windows.

### Statistics

Statistical analysis was performed in Matlab (MATLAB R2018a, MathWorks, USA) and in Excel (Excel 2016). Significance between data sets was tested by one-way or two-way ANOVA, as indicated; significance is given as p value (*p ≤ 0.05; **p ≤ 0.01; ***p ≤ 0.001).

