## Supplemental Figures for "Coordinated electrical and chemical signaling between two neurons orchestrates switching of motor states"

### Supplemental Material

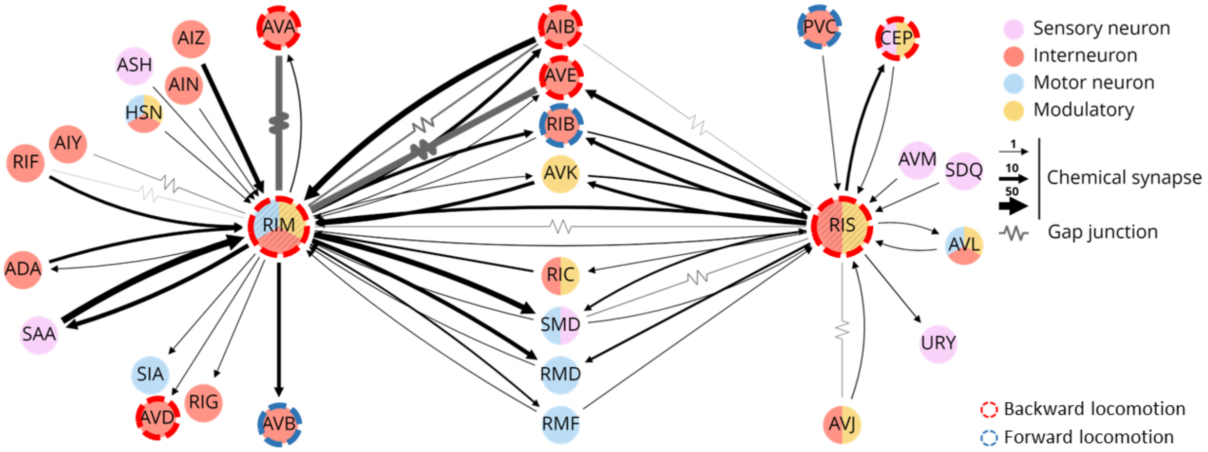


**Figure S1: Neuronal networks involving RIS and RIM neurons**

RIS and RIM connectome with neuron types as indicated by color. Black arrows represent chemical synapses and grey lines gap junctions. Thickness indicates abundance of connections. Neurons known to promote forward or backward locomotion are additionally indicated by dashed blue or red circles, respectively. Modified from nemanode.org (based on Refs. [24, 25]).

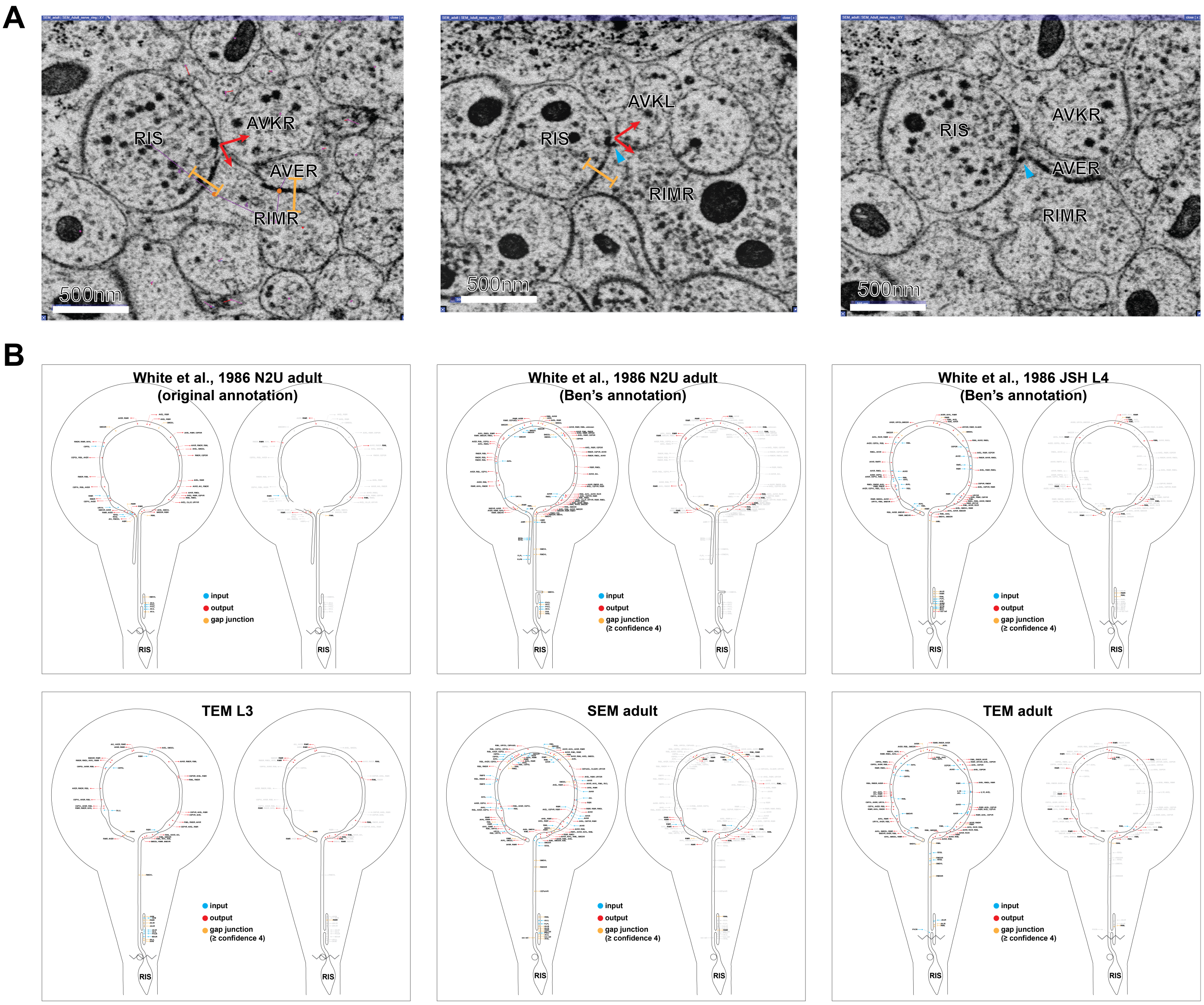


**Figure S2: RIS morphology and electron microscopy-based wiring diagrams**

A) Example electron micrographs showing aspects of RIS and RIM connectivity. Cellular processes / axons are identified. Yellow lines indicate gap junctions, red arrows label chemical synaptic output. Blue arrowhead marks potential postsynaptic density. Scale bar is 500 nm.

B) Annotations of synapses are placed on schematic showing RIS morphology. The left panels show all confidently annotated chemical synapses and gap junctions from the White et al original annotation, my reannotation, and the Witvliet et al paper. The right panels simply highlight RIS-RIM connections. Top three panels: Annotations of [24] are indicated, as well as our own annotations of the White data set ("Ben's annotation"). Lower panels: RIS synapses deduced from three reconstructions (left to right: L3 animal, analyzed by TEM on individual thin sections; adult animal, imaged by SEM on tape-affixed thin sections; adult animal imaged by TEM on individual thin sections. Chemical input (blue) and output (red) synapses, as well as electrical synapses (yellow) are labeled.

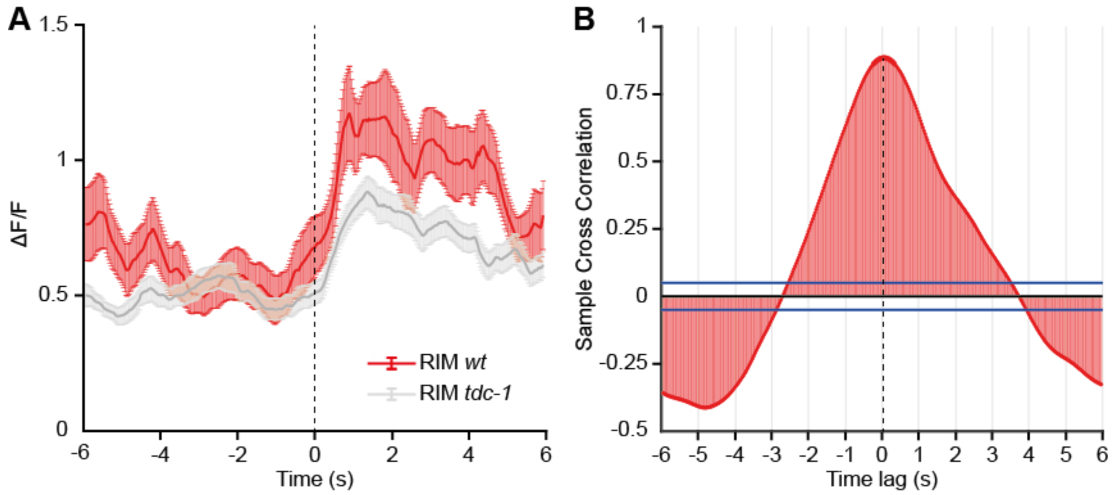


**Figure S3: RIM $Ca^{2+}$ compared in wild type and *tdc-1(n3419)* mutants.**

A) $Ca^{2+}$-signals of RIM in free moving wild type (red, N = 9 animals, n=20 events) and *tdc-1* mutant (grey; N = 15, n=53) aligned to locomotion stop. Mean ± SEM of 6 s time windows before and after locomotion stop.

B) $Ca^{2+}$-signals of RIM in wild type and *tdc-1* mutants show significant cross correlation and no time lag (N = 15, n = 53 for *tdc-1* and N=9, n = 20 for WT, Pearson's r = 0.89). Blue bars indicate 95% confidence bounds.

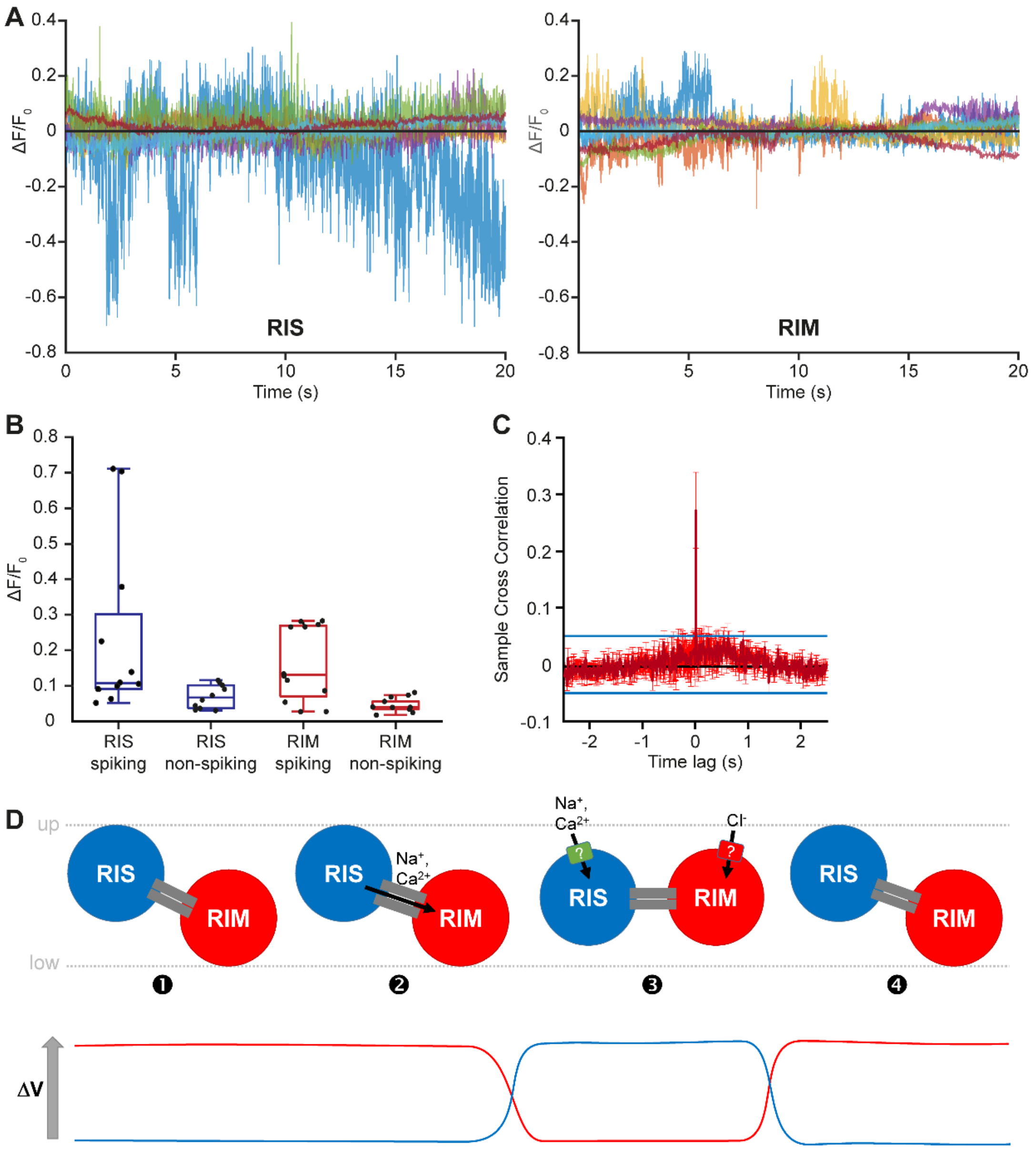


**Figure S4: Signal properties of RIS and RIM during dual voltage imaging**

(A) All spiking traces of RIS (left panel) and RIM (right panel).

(B) Amplitudes of spiking and non-spiking activity in 5 s windows for RIS (blue) and RIM (red), used for cross correlation analysis.

(C) Cross correlation analysis of RIS and RIM GFP signals of spiking animals. Mean ± SEM of 5 s time windows; blue lines indicate 95% confidence bounds.

(D) Model for how oscillating electrical activity may arise between RIS and RIM (and vice versa, not shown). Initially (1), RIS depolarizes and the (rectifying) gap junction (grey bars) opens (2), allowing current to leave RIS. As a consequence, RIS drops and RIM rises in voltage level. The fluorescent signals are indicated below. (3) the Gap junctions close, and unknown (ligand gated) ion channels in RIS and RIM (depolarizing or hyperpolarizing, as indicated by positive and negative ions) open to restore the initial voltage levels (4). The nature and identity of these channels is unknown, but some of these activities may be due to tyraminergic $Cl^-$ channels, possibly in (auto-) feedback loops between the two neurons (speculative). The opposing situation, which was also found experimentally, i.e. RIM being in an up- and RIS in a down-state, is not shown, for simplicity, but would be analogous, with the gap junction rectifying in the opposite direction. Note that the indicated fluorescent voltage signals do not report on absolute voltage in the two neurons (which is not known), but to relative changes over time.

RIS gap junctions 2022

White et al., 1986
N2U adult

| Cell | # gap junctions |
|---|---|
| AVJL | 2 |
| RIML | 1 |
| RIMR | 1 |
| SMDVL | 1 |
| SMDDL | 1 |
| SMDDR | 1 |
| AIBR | 1 |

SEM adult

| | | # gap junctions confidence level | | | | |
|---|---|---|---|---|---|---|
| | | 5 | 4 | 3 | 2 | 1 |
| Cell | RIMR | 1 | 3 | 2 | | |
| | RIML | 1 | 1 | 2 | 1 | |
| | exc cell | 1 | 2 | | | |
| | AVJR | 1 | 1 | | | |
| | AVEL | 1 | 1 | | | |
| | AVJL | 1 | | | | |
| | SMDVL | | 2 | 2 | 1 | |
| | RIBL | | 2 | | | |
| | RIBR | | 1 | 2 | 1 | |
| | SMDDR | | 1 | 1 | 3 | |
| | SMDDL | | 1 | 1 | 1 | |
| | RMDDR | | 1 | | | |
| | RMEL | | 1 | | | |
| | CEPshVR | | 1 | | | |
| | RMDL | | | 4 | 2 | |
| | RMDR | | | 2 | 1 | |
| | RMED | | | 2 | | |
| | SMDVR | | | 1 | | |
| | RMER | | | 1 | | |
| | AVBL | | | 1 | | |
| | AVBR | | | 1 | | |
| | RICL | | | 1 | | |
| | AIBL | | | 1 | | |
| | CEPVL | | | 1 | | |
| | AVER | | | | 1 | |
| | CEPDR | | | | 1 | |
| | URYVR | | | | 1 | |
| | RIVL | | | | 1 | |
| | SMBVR | | | | 1 | |
| | RIH | | | | 1 | |

TEM adult

| | | # gap junctions confidence level | | | | |
|---|---|---|---|---|---|---|
| | | 5 | 4 | 3 | 2 | 1 |
| Cell | RIML | 2 | | | 2 | |
| | AVJR | 1 | | | | |
| | SMDVL | | 2 | 1 | 1 | |
| | RMDDR | | 2 | | | |
| | AVEL | | 1 | 1 | | |
| | AVER | | 1 | | | |
| | RIBR | | 1 | | | |
| | RIMR | | | 1 | 2 | |
| | CEPVR | | | 1 | | |
| | RMDL | | | 1 | | |
| | CEPVL | | | 1 | | |
| | SMDDL | | | 1 | | |
| | CEPshVR | | | 1 | | |
| | SMDDR | | | | 2 | |

TEM L3

| | | # gap junctions confidence level | | | | |
|---|---|---|---|---|---|---|
| | | 5 | 4 | 3 | 2 | 1 |
| Cell | AVJR | 1 | 1 | | | |
| | RIMR | | 2 | 1 | | |
| | AVM | | 2 | | | |
| | AVJL | | 1 | | | |
| | RMDVL | | 1 | | | |
| | SMDDR | | | 2 | | |
| | RIML | | | 2 | | |
| | CEPDL | | | 1 | | |
| | CEPVL | | | 1 | | |
| | AVBL | | | 1 | | |
| | SMDVR | | | 1 | | |
| | AIYL | | | 1 | | |
| | RMED | | | 1 | | |
| | RMEV | | | | 2 | |

N2U adult from White et al., 1986
Ben's annotation

| | | # gap junctions confidence level | | | | |
|---|---|---|---|---|---|---|
| | | 5 | 4 | 3 | 2 | 1 |
| Cell | AVEL | 2 | | | | |
| | SMDVL | 1 | 2 | | | |
| | AIBR | 1 | 1 | | | |
| | SMDDR | 1 | | 2 | 1 | |
| | RIML | 1 | | | 1 | |
| | AVJL | 1 | | | | |
| | RMDVL | 1 | | | | |
| | DVA | 1 | | | | |
| | SMDDL | | 2 | | | |
| | RIMR | | 1 | | | |
| | CEPDR | | 1 | | | |
| | SMDVR | | | 3 | | |
| | FLPR | | | 1 | | |
| | AIBL | | | 1 | | |
| | RIBR | | | 1 | | |
| | RICR | | | 1 | | |
| | RMDDL | | | 1 | | |
| | SIAVR | | | 1 | | |
| | AVM | | | | 1 | |
| | CEPDL | | | | 1 | |
| | CEPVL | | | | 1 | |
| | RIBL | | | | 1 | |
| | RMEV | | | | 1 | |
| | SMDDR | | | | 1 | |
| | URYVL | | | | 1 | |

JSH L4 from White et al., 1986
Ben's annotation

| | | # gap junctions confidence level | | | | |
|---|---|---|---|---|---|---|
| | | 5 | 4 | 3 | 2 | 1 |
| Cell | RIMR | 1 | 2 | 1 | | |
| | AVJR | 1 | 1 | | | |
| | RIML | 1 | | 2 | | |
| | SMDDL | 1 | | 1 | | |
| | AVJL | 1 | | | | |
| | AIBL | 1 | | | | |
| | SMDDR | | 1 | | | |
| | RIBR | | | 1 | | |
| | AVER | | | | 1 | |
| | RMED | | | | 1 | |

### Supplemental Table 1

Summary of annotations of gap junctions observed between RIS and RIM. Where indicated, data were included from Ref. 24 (White et al. 1986). Other data are from Ref. 25 and this work.
